# Immigration delays but does not prevent adaptation following environmental change: experimental evidence

**DOI:** 10.1101/2023.08.31.555763

**Authors:** Lily F. Durkee, Laure Olazcuaga, Brett A. Melbourne, Ruth A. Hufbauer

**Author notes:** Journal of Evolutionary Biology* special issue: Eco-evolutionary dynamics in changing environments.

## Abstract

In today’s rapidly changing world, it is critical to examine how animal populations will respond to severe environmental change. Following events such as pollution or deforestation that cause populations to decline, extinction will occur unless populations can adapt in response to natural selection, a process called evolutionary rescue. Theory predicts that immigration can delay extinction and provide novel genetic material that can prevent inbreeding depression and facilitate adaptation. However, when potential source populations have not experienced the new environment before (*i.e.,* are naive), immigration can counteract selection and constrain adaptation. This study evaluated the effects of immigration of naive individuals on evolutionary rescue using the red flour beetle, *Tribolium castaneum,* as a model system. Small populations were exposed to a challenging environment, and three immigration rates (zero, one, or five migrants per generation) were implemented with migrants from a benign environment. Following an initial decline in population size across all treatments, populations receiving no immigration gained a higher growth rate one generation earlier than those with immigration, illustrating the constraining effects of immigration on adaptation. After seven generations, a reciprocal transplant experiment found evidence for adaptation regardless of immigration rate. Thus, while the immigration of naive individuals briefly delayed adaptation, it did not increase extinction risk or prevent adaptation following environmental change.

## 1 Introduction

A critical goal of conservation biology is to establish how to effectively manage populations in human-altered habitats. Environmental change such as pollution (Loria *et al*., 2019), deforestation (Foster *et al*., 2021), and fragmentation (Cheptou *et al*., 2017) can reduce habitat quality and connectivity, which can decrease the average fitness of residing populations. If fitness drops below the replacement rate, populations will decline in size and will approach extinction if the downward trend persists. Immigration, either through maintenance of dispersal corridors or translocations, is one approach readily available to conservation practitioners to support declining populations, particularly in fragmented landscapes (Frankham *et al*., 2019).

Theoretical work within the conservation literature recommends an immigration rate of one to ten migrants per generation for isolated populations (Mills & Allendorf, 1996; Frankham *et al*., 2019). More specifically, theory suggests a rate of one migrant per generation to offset population divergence and loss of heterozygosity that occurs through drift (Mills & Allendorf, 1996), and five migrants per generation to prevent damaging inbreeding depression and maintain adaptive genetic variation in changing environments (Frankham *et al*., 2019). Ideally, immigration can occur consistently through dispersal corridors, and if translocations are needed, consistent immigration should be sustained. However, these recommendations have been tested only partially. Laboratory experiments have demonstrated the benefits of immigration when comparing isolated populations to populations that received a single immigration event under scenarios of inbreeding (Spielman & Frankham, 1992) or low diversity populations facing environmental change (Hufbauer *et al*., 2015). Only one study we know of, Newman & Tallmon (2001), tested immigration that occurred every generation. They assessed rates of one or 2.5 migrants per generation in recently isolated mustard plant populations, and found that the rate of one migrant per generation was sufficient to maintain genetic variation and decrease inbreeding when compared to isolated populations (Newman & Tallmon, 2001). In this era of global change, it is important to also test these recommendations under more severe environmental conditions.

A separate but related body of literature focuses on how populations declining due to exposure to challenging environments can avoid extinction through adaptive evolution, a process called evolutionary rescue (Gomulkiewicz & Holt, 1995; Carlson *et al*., 2014). Examples include adaptation in weed or insect populations to pesticides, antibiotic resistance in bacteria, or invertebrate adaptation to metal pollution in streams (Bell, 2017). Experiments show that immigration or admixture can enhance the probability of evolutionary rescue if populations are exposed to a challenging environment (Hufbauer *et al*., 2015; Durkee *et al*., 2023). Furthermore, theoretical studies show that the probability of rescue can increase with immigration, primarily due to the added genetic variation, which increases the chance of beneficial alleles being present (Tomasini & Peischl, 2020; Czuppon *et al*., 2021). To contrast, other theoretical studies show that gene flow from maladapted immigrants can limit adaptation by swamping adaptive alleles in the recipient population, reducing fitness (*i.e.,* outbreeding depression) (Slatkin, 1987; Lenormand, 2002; Brady *et al*., 2019).

To reduce the likelihood of outbreeding depression, using a source population from a habitat similar to the recipient population (*i.e.,* habitat matching) is recommended by models (Edelaar & Bolnick, 2012) and in the context of conservation (Frankham *et al*., 2019). However, this is often not possible, particularly under scenarios of novel environmental change (Aitken & Whitlock, 2013) or when the species of concern is rare (Frankham *et al*., 2019). Even when migrants are not adapted to their new habitat, for example when individuals from a geographically distant population (*e.g.,* Hedrick & Fredrickson, 2010) or from captivity (*e.g.,* Crone *et al.,* 2007) are used, every-generation immigration of one to five individuals is meant to be low enough to minimize outbreeding depression while still maintaining genetic diversity and adaptive potential of populations (Newman & Tallmon, 2001; Frankham *et al*., 2019). However, to our knowledge, no study has compared the recommended immigration rates of one and five migrants per generation and their impacts on adaptation in the context of severe environmental change.

Here, we evaluated how immigration rates recommended in the conservation literature (Frankham *et al*., 2019) impacted evolutionary rescue (Carlson *et al*., 2014). To do this, we first created experimental populations of red flour beetles (*Tribolium castaneum*) from a genetically admixed stock population (Durkee *et al.,* 2023). We then introduced them to a challenging environment to mimic a situation where a large, diverse population experiences sudden habitat change that decreases population size and reduces habitat quality, for example when forest are logged (Hillers *et al.,* 2008). Each experimental population received one of three immigration treatments (zero, one, or five individuals every generation), and we observed the effects of immigration on population persistence and growth. In habitats like these that have limited space and degraded resources, population recovery may be constrained by density-dependent processes like competition for food and space (Osmond & de Mazancourt, 2013; Nordstrom *et al*., 2023). Thus, we evaluated both density-dependent and density-independent population growth through time to tease apart the impacts of negative density dependence and adaptation. Our experiment was designed to address the following question: In populations experiencing severe environmental change, how do different immigration rates affect *(i)* population size and growth rates over time, *(ii)* the timing of evolutionary rescue, and *(iii)* adaptation to the challenging environment?

## 2 Methods

To evaluate the question posed in the introduction, we carried out an experiment where we subjected replicated, independent populations to a challenging environment that caused them to decline in size. This environment, therefore, would have led to extinction unless the populations were able to adapt sufficiently to the challenging conditions. Hereafter, we refer to this as the rescue experiment (Figure 1a-b).

**Figure 1.**
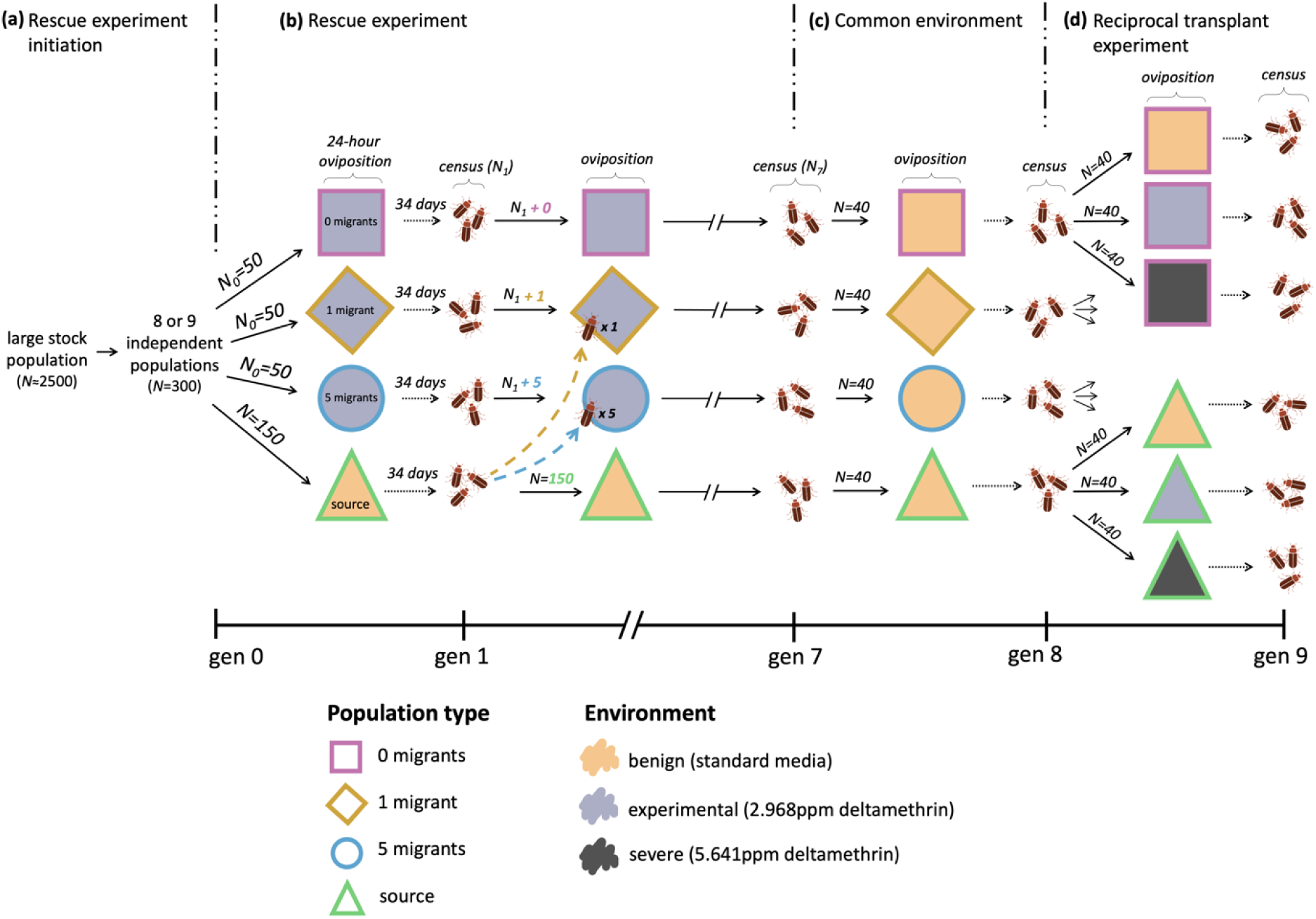
Experimental design for the rescue and reciprocal transplant experiments. **a)** We began with a large stock population (*N*≈2500), from which eight or nine independent populations were initiated (*N*=300). Each of these populations was split into one source population, initiated and then maintained at *N*=150 individuals, and three experimental populations, initiated with *N*_0_=50 individuals with population size allowed to fluctuate thereafter. The source populations were maintained on a benign environment (tan shading), and the experimental populations were maintained in a challenging environment containing the insecticide deltamethrin (light gray shading). **b)** In the rescue experiment, census events (gens 1 & 7 shown) are indicated with three cartoon beetles, and census occurred every generation on day 35, following a 24-hour oviposition period and 34 days of development (indicated with dotted arrows). Immigration from the source to the experimental populations (dashed arrows) occurred after each census event at the start of the oviposition period, and three treatments were implemented: 0 migrants (purple square outlines), 1 migrant (yellow diamond outlines), and 5 migrants per generation (blue circle outlines). **c)** All extant populations were placed on a benign, common environment for a single generation following the census in generation seven at a density of *N*=40, with as many replicates created as possible. **d)** In the reciprocal transplant experiment, all populations were split into groups of 40 and placed simultaneously and at constant densities on three different environments: benign, challenging, and severe (dark gray shading). This whole procedure **(a-d)** was repeated across three temporal blocks, with eight replicates in blocks 1 and 2 and nine replicates in block 3.

### Rescue experiment initiation and propagation

We created a genetically diverse *Tribolium castaneum* stock population comprised of lineages described in Durkee *et al*. (2023). The stock population had been reared on standard media (19:1 wheat flour to brewer’s yeast) and in standard environmental conditions (31°C and 50-80% humidity) in 4x4x6cm patches for nine, 35-day non-overlapping generations prior to the start of the rescue experiment. To start, we created 25 independent populations from the stock population of 300 individuals each (Figure 1a). These populations were used to initiate replicated experimental populations and maintain populations to serve as a source of migrants (hereafter, source populations). These populations were divided at the start of the rescue experiment, and therefore represent newly isolated populations with no history of divergence, similar to Newman & Tallmon (2001).

#### Experimental populations

were each initiated with 50 adult beetles, 36 days old, and population sizes were allowed to fluctuate naturally in subsequent generations. Each population was placed onto a single patch of challenging environment created using media containing the pyrethroid insecticide deltamethrin (DeltaDust, Bayer). This event represents a sudden environmental change that reduced fitness and created replicated, isolated populations. The experimental patches each contained a concentration of deltamethrin that a dose-response curve created from preliminary experiments (Figure S1) indicated would decrease the population growth rate to approximately 0.40 from a typical value of around 2. Without adaptation, this reduced growth rate should have led to extinction in 10 generations (based on simulations with the stochastic Ricker model developed for *T. castaneum* by Melbourne & Hastings, 2008). The experimental populations (*N*=50) were initiated in 25 groups of three, each group from one of the original 25 independent populations (Figure 1a).

#### Source populations

were initiated with 150 adults, 36 days old, and each served as a source of migrants for a group of three experimental populations (Figure 1a). Thus, 25 source populations were initiated in total, each from one of the original 25 populations. We intended for the sources to represent larger populations, and so we used the large population size (*N*=150) from Hufbauer *et al*. (2015). Sources were maintained at 150 adults every generation regardless of the number of offspring produced, to approximate populations with a carrying capacity of 150 individuals. Source populations were each maintained on a single patch containing standard dry media, a benign environment for *T. castaneum*. Thus, these individuals were not exposed to the challenging environment present in the experimental patches. Hereafter, we refer to these individuals as naive migrants, to refer to both evolutionary and behavioral naivety.

The adult beetles in the experimental and source patches were given 24 hours to mate and oviposit. Adults were then were discarded, and the eggs developed for an additional 34 days. The first census event (*N*_1_, Figure 1b) occurred on day 35. Subsequently, one of three immigration treatments was applied to each experimental population in every group: zero, one, or five migrants. The migrants were moved from the source to each experimental patch manually using a paintbrush. This occurred at the start of the following oviposition period. This procedure (immigration of zero, one, or five migrants, 24-hour oviposition, removal of adults, 34-day development period, census) repeated each subsequent generation. The experiment was terminated after seven generations because nearly all the experimental populations had either gone extinct or their growth rates and population sizes had increased, indicative of evolutionary rescue (see Figure 2).

**Figure 2.**
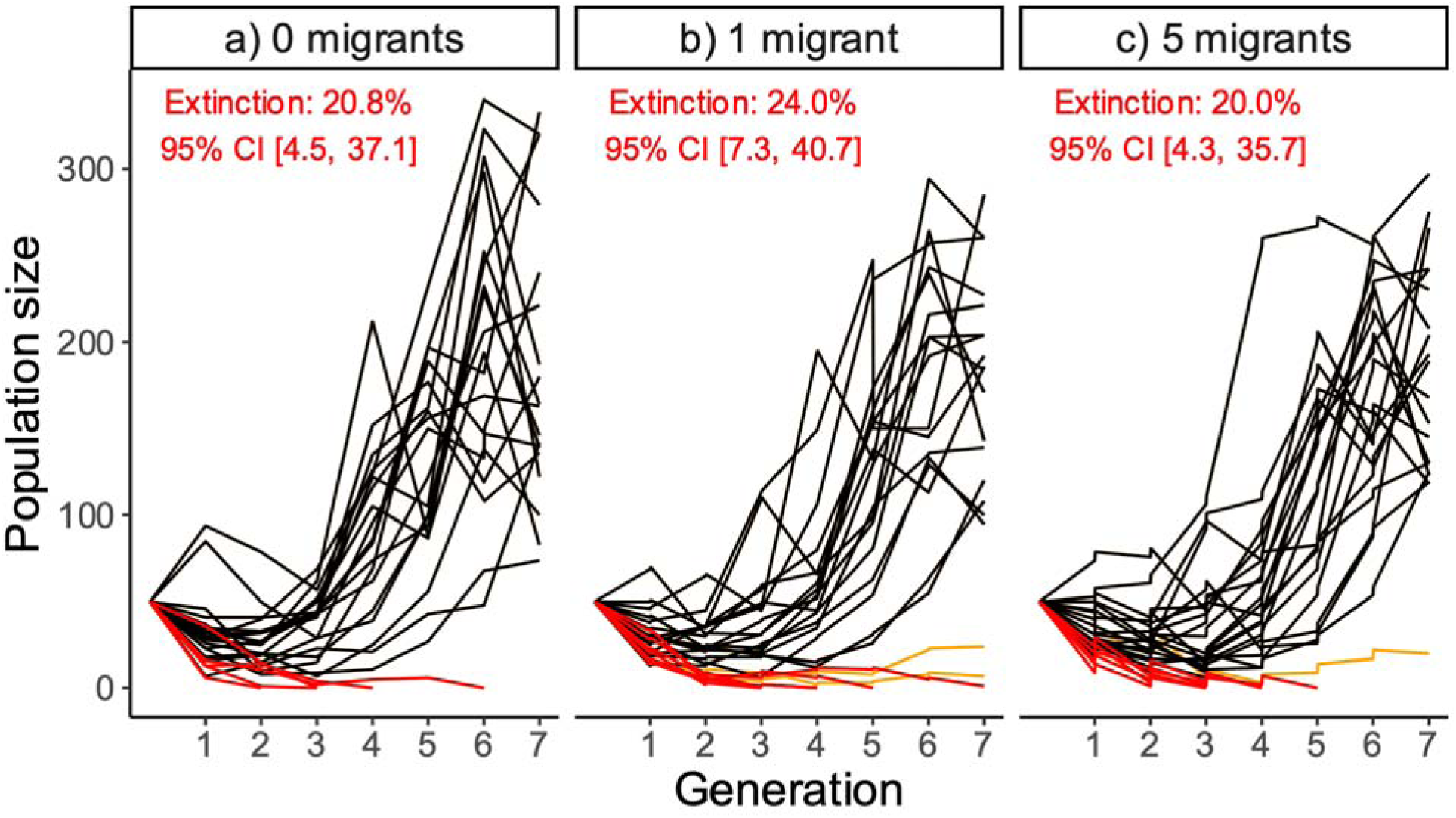
Population size through time in discrete generations for the populations receiving 0, 1, or 5 migrants per generation during the rescue experiment. Population sizes represent the number of adult beetles present at census (day 35 each generation). Each black line represents a rescued population, each orange line represents an extant population that was not considered rescued, and each red line represents a population that went extinct or had a population size of one individual in generation seven. Immigration events are indicated with instantaneous increases in population size, which occurred at the start of the first six generations. The percent of populations that went extinct and simple confidence intervals around each percent are provided.

In summary, there were 25 groups, each consisting of one source and three experimental populations (one of each treatment). Thus, there were 25 replicates of each treatment, which we split across three temporal blocks (eight groups in blocks 1 and 2, nine in block 3; Figure 1a). Within each group, treatments were set up sequentially, and propagated for the first four generations in the following order to minimize opportunities for experimental error: 1-migrant, 5-migrants, and then the 0-migrant populations. As this was not random, we tested if this order mattered in a follow-up experiment and found no significant effect of treatment order on the growth rate the following generation (Figure S2). During generations 4 through 7, the experimental protocol was well established, and so we randomized the order of set-up.

### Source population dispersal arrays

The immigration recommendations we followed (one or five migrants per generation) are based on either natural dispersal or translocation. Thus, in order to approximate natural dispersal and make our results applicable to either case, we used migrants with higher demonstrated dispersal ability to represent individuals that would be more likely to leave the source patch and move to a new patch in the absence of translocation (Edelaar & Bolnick, 2012; Cote *et al*., 2022). Dispersal arrays were created to select dispersing individuals. On day 34 during generations 1-6, habitat patches housing source populations were connected to 3-patch dispersal arrays connected with 2mm holes, creating a 4-patch dispersal array. Hereafter, we refer to the original source patch as patch 1 and the subsequent patches, in order, as patches 2-4, which contained media that had been previously used to rear beetles for 35 days to keep resource availability consistent throughout the array. Adult beetles were given 24 hours to disperse out of patch 1. On day 35, all adults were censused in each of the four patches of the dispersal array (Figure S3). Individuals that dispersed to patches 2-4 were used as migrants (hereafter called dispersing migrants) with preference given to individuals that dispersed to the furthest patch (*i.e.,* individuals were first chosen from patch 4, then patch 3, then 2).

In order to maintain the population size of the source populations at 150 to start every generation, growth rates needed to be above the replacement rate. However, growth rates were sometimes lower than expected, which likely occurred due to effects of demographic stochasticity, which strongly influences the *T. castaneum* system (Melbourne & Hastings, 2008). When this happened, populations were supplemented with individuals from the stock population at the start of the oviposition period, after the dispersal period and immigration to the experimental populations occurred. After the first generation, two source populations were initialized with 110 individuals due to limited availability of stock individuals.

### Common environment and reciprocal transplant experiment

After seven generations, we terminated the rescue experiment, and all extant populations (58 experimental and 25 source populations) were placed at a constant density of 40 individuals on standard media for a single generation (hereafter, “common environment” generation; Figure 1c). All adults were kept within their experimental populations, *i.e.,* no mixing occurred within or across treatments. In *T. castaneum,* maternal environmental effects can strongly affect growth rate (Hufbauer *et al.,* 2015; Van Allen & Rudolf, 2013), and so this generation where the environment and density were kept the same for all populations served to standardize maternal effects for the subsequent reciprocal transplant experiment. With only a single generation, we do not standardize grand-maternal effects. However, inference from multigenerational studies suggests that grand maternal effects are relatively weak in *T. castaneum* populations (Van Allen & Rudolf, 2013; Hufbauer *et al.,* 2015).

In the ninth generation, a reciprocal transplant experiment was completed to assess adaptation to the challenging environment and potential costs to adaptation in the source environment (Figure 1d). We divided individuals from each population into three groups and randomly assigned them to one of three environments: standard media (0ppm, hereafter “source environment”), media containing the concentration of deltamethrin present in the challenging environment of the rescue experiment (2.968ppm, hereafter “experimental environment”), and media containing a higher concentration of deltamethrin (5.641ppm, hereafter “severe environment”). The dose-response curve (Figure S1) indicated that the severe environment would reduce the growth rate of naive populations to near zero. Each replicate patch of these three environmental treatments received 40 beetles, which were allowed 24 hours for oviposition, and 34 additional days for development prior to census. Replication within population depended on the number of available individuals following the common environment generation (see Figure S4).

### Data analysis

All statistical analyses were performed in R version 4.1.2 (R Core Team, 2021). Population size at census, which occurred on day 35 during generations 1-7, was used to assess population dynamics throughout the rescue experiment. Populations were considered rescued if they grew to a population size greater than 50, and maintained a growth rate greater than 1 (the replacement rate) for at least two of the final four generations. This allowed for decreases in population size due to negative density dependence. Extinction risk and survival times were quantified using a survival analysis with the *ggsurvfit* function in the *survival* package (Therneau 2021). Extinct populations were defined as those with zero adults at census during the first six generations or one adult by generation seven. Overall extinction rates were calculated as raw percentages with confidence intervals around each percentage for each treatment.

Population growth rate was calculated as the mean number of surviving adult offspring per individual in the population, *i.e.,* population size at census each generation (*N*_t_) divided by the number of parents that produced those individuals (parent population size, *N*_t-1_), or *N*_t_/*N*_t-1_. We evaluated both density-dependent and density-independent growth rates through time using a linear mixed model with the *lme4* package (Bates *et al*., 2015). We included immigration treatment (0 migrants, 1 migrant, or 5 migrants per generation), time (*t*) in generations (from 1 to 7 generations, categorical), and parent population size (*N*_t-1_, integers) as fixed effects, and temporal block (block 1, block 2, block 3), source population (nested within block), and population identity (nested within source population) as random effects. Growth rate data were *log(x+1)* transformed to help satisfy the equal variance assumption. Diagnostic plots were visualized for all models using the *DHARMa* package (Hartig 2022).

From this model, we generated estimates of both density-dependent and density-independent population growth rates using the *predict* function with different values for the parent population size (*N*_t-1_) each generation. To estimate density-dependent growth rates, we allowed the values for *N*_t-1_ to vary for each generation and treatment combination, matching the observed mean population sizes at the start of each generation. This estimate was used to describe the population growth rate data and quantify the effects of immigration treatment in a finite habitat by considering the constraints of competition for resources and cannibalism that affected the populations in our experiment (Pointer *et al*., 2021). To estimate density-independent growth rates from the fitted model, we set *N*_t-1_ to zero for every generation and treatment, which is equivalent to estimating the y-intercepts when plotting model-estimated growth rate over *N*_t-1_ (Figure S5). These density-independent estimates allowed us to track adaptation to the environment by revealing the change in intrinsic fitness of individuals in the population unfettered by the effects of density. We expected the density-independent growth rates to be higher than the density-dependent estimates due to the known negative density dependence in *T. castaneum* (Melbourne & Hastings 2008). Confidence intervals for both density-dependent and density-independent growth rate estimates were estimated by parametric bootstrap (percentile method) using the *bootMer* function with 5000 simulations in the *lme4* package (Bates *et al*., 2015). Comparisons between treatments were made by estimating confidence intervals by parametric bootstrap (percentile method) for the difference in means between treatments within each generation using the *simpleboot* and *boot* packages with 5000 simulations (Peng, 2019; Canty & Ripley, 2022).

The timing of evolutionary rescue was assessed in each treatment group using a Generalized Additive Mixed (GAM) model (Wood, 2017) similar to Olazcuaga *et al*. (2023). Only rescued populations were considered in this analysis. We modeled the population size of rescued populations using a negative binomial distribution to account for overdispersion using the package *mgcv* (Wood, 2004, 2011). To allow for the relationship between the population size and time to be nonlinear, we modeled population size as a smooth function of generation with the number of knots set to 4, which was chosen to maximize the adjusted R-squared value. Explanatory variables also included immigration treatment and the interaction of treatment and generation as fixed effects, allowing for a smooth function for each treatment. The full model included population identity and temporal blocks as random effects. The best-fit model was chosen by comparing to reduced models using the *compareML* function from the *itsadug* package (van Rij *et al*., 2022).

We used data from the reciprocal transplant experiment to further evaluate adaptation to the experimental environment, as well as potential fitness costs of adaptation in a benign environment. Growth rate in the reciprocal transplant experiment was square-root transformed and used as the response variable in a linear mixed effects model that included population type (source, 0-migrant, 1-migrant and 5-migrant), test environment (source, experimental, severe), and the interaction between population type and test environment as fixed effects. Population identity was included as a random effect nested within temporal block. Comparisons within each environment and among immigration treatments were made using 95% bootstrapped confidence intervals (percentile method) for the difference in means, which were generated from 5000 iterations.

## 3 Results

### Population size and extinction

During the rescue experiment, a majority of populations experienced evolutionary rescue (Figure 2, black lines), and extinction probability was similar for all three treatments (Figure 2, red lines). The difference in the extinction risk was statistically similar across treatments throughout the experiment (Figure S6). Additionally, there was no clear difference in survival time among treatments 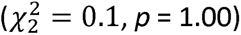.

### Growth rate through time

Density-dependent growth rates increased through generation four or five, then decreased again as population density increased (Figure 3a, Table S1). Density-independent growth rates were below or close to the replacement rate (growth rate of 1) in generations 1-3 (Figure 3b; confidence intervals overlap with dashed line). This changed in generation four, when the 0-migrant treatment had a mean density-independent growth rate much larger than the replacement rate, illustrating rapid adaptation within those populations (Figure 3b). In generation four, density-independent growth rate in the 0-migrant populations was on average 0.808 (95% CI [0.207, 1.453]) higher than the 1-migrant populations and 0.969 [0.478, 1.573] higher than the 5-migrant populations. By generation five, all treatments gained and then maintained a mean density-independent growth rate greater than 1, demonstrating adaptation to the challenging environment. The comparison between the density-independent and density-dependent estimates of growth rates suggest that the decrease in the latter by generation seven was likely due to the effects of negative density dependence, which increased as populations grew.

**Figure 3.**
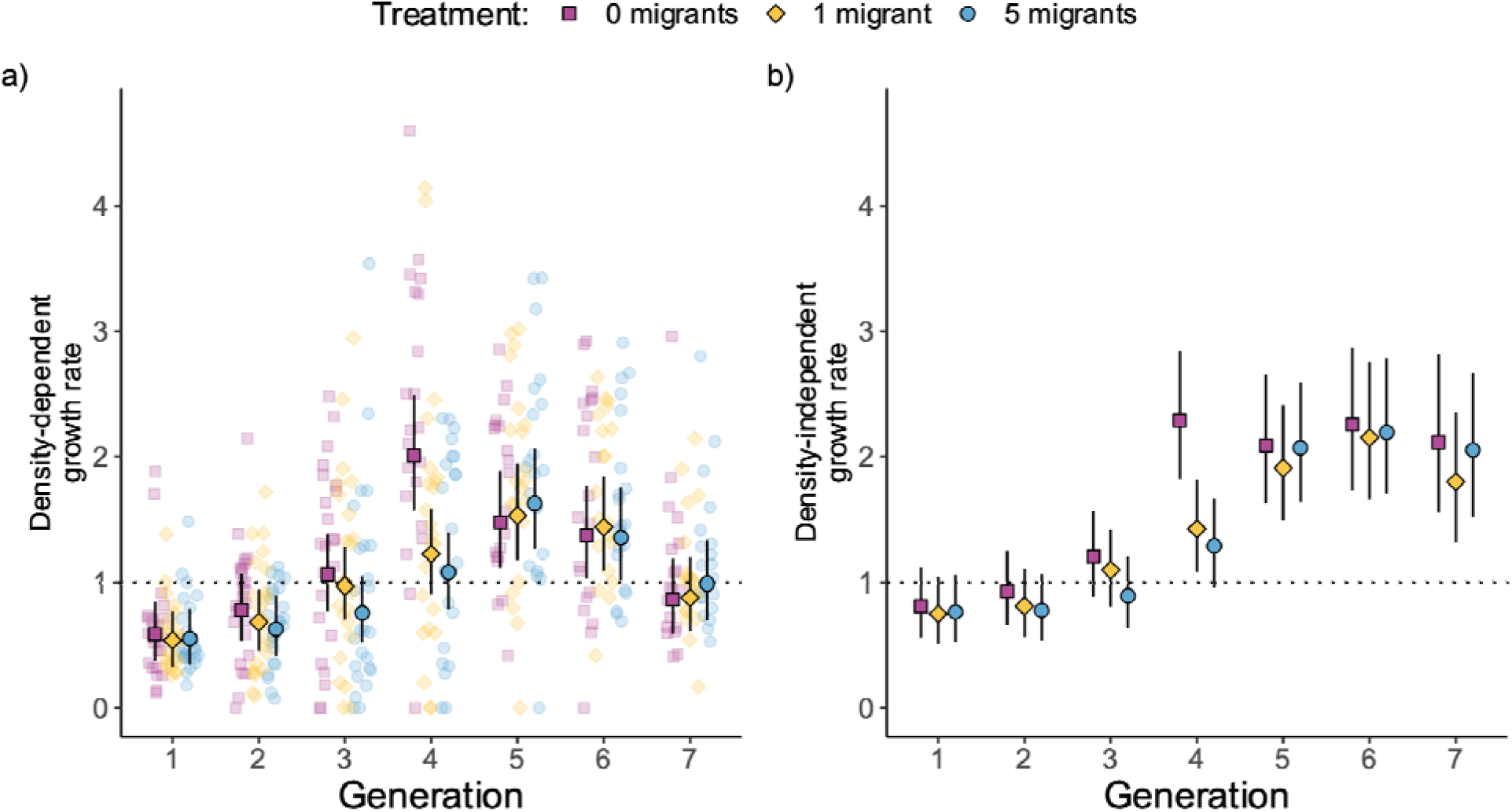
Density-dependent growth rate (a) and density-independent growth rate (b) throughout the rescue experiment. Each point outlined in black is the predicted mean for each treatment (purple square = 0 migrants, yellow diamond = 1 migrant per generation, and blue circle = 5 migrants per generation). Error bars are bootstrapped 95% confidence intervals. Points without black outlines are raw growth rate values, and each point represents one population. The dotted line represents a growth rate of 1, the replacement rate. Two estimates for growth rate are presented: a) the density-dependent estimates are predictions from the linear mixed model for each generation and treatment combination, used to incorporate the effects of negative density-dependence and provide estimates that describe the raw data (shown), and b) the density-independent estimates (i.e., predictions using N_t-1_ = 0), showing adaptation over time.

### Timing of evolutionary rescue

Among the rescued populations only (black lines in Figure 2), the analysis of population size with a GAM model revealed a shallow U-shaped curve (Figure 4). The estimated curves from the best-fit model, which included treatment and the interaction of treatment and generation as fixed effects and population identity as a random effect, are shown in Figure 4. The interaction between immigration treatment and generation was significant (, *p* < 0.0001), with the 0-migrant populations growing slightly faster than populations receiving migrants (Figure 4). The 1-migrant and 5-migrant treatments maintained similar population trajectories throughout the experiment.

**Figure 4.**
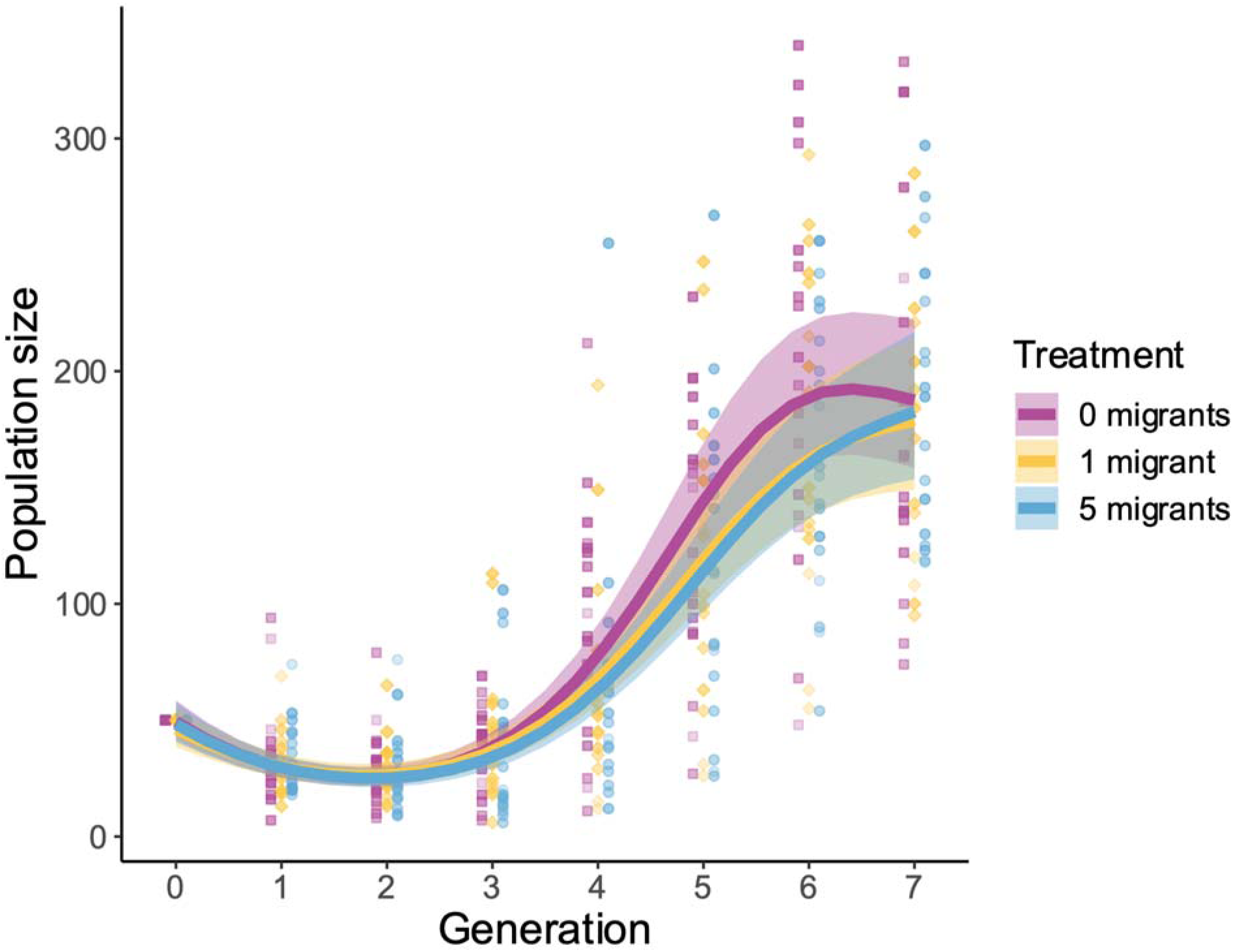
Evolutionary rescue analysis. Population size over time (in generations) of rescued populations receiving 0 migrants (purple line, squares), 1 migrant (yellow line, diamonds), or 5 migrants (blue line, circles) per generation estimated by the best-fit GAM model. Each point represents one population, which are staggered to reduce overlap. The lines are estimates of mean population size for the three immigration treatments, and the shaded region around each line is the 95% confidence interval given by the GAM model.

### Adaptation and associated costs

Growth rate in the reciprocal transplant experiment was influenced by an interaction between environment and population type (F_6, 168.04_ = 25.16, *p* < 0.0001), in a pattern showing that adaptation to the pesticide was linked to reduced fitness in the benign source environment (Figure 5). In the benign environment, the experimental populations had lower growth rates on average compared to the source populations (confidence intervals are negative and do not overlap with zero: 95% CI_0-migrant – source_ [-0.855, -1.628]; 95% CI_1-migrant – source_ [-0.911, -1.748]; 95% CI_5-migrant – source_ [-0.641, -1.482]), and a higher growth rate in the severe habitat (confidence intervals are positive and do not overlap with zero: 95% CI_0-migrant – source_ [0.442, 0.171]; 95% CI_1-migrant – source_ [0.424, 0.210]; 95% CI_5-migrant – source_ [0.464, 0.216]). Additionally, the 5-migrant populations performed marginally better than the 0-migrant populations in the experimental environment (95% CI_0-migrant – 5-migrant_ [-0.0675, 0.572]).

**Figure 5.**
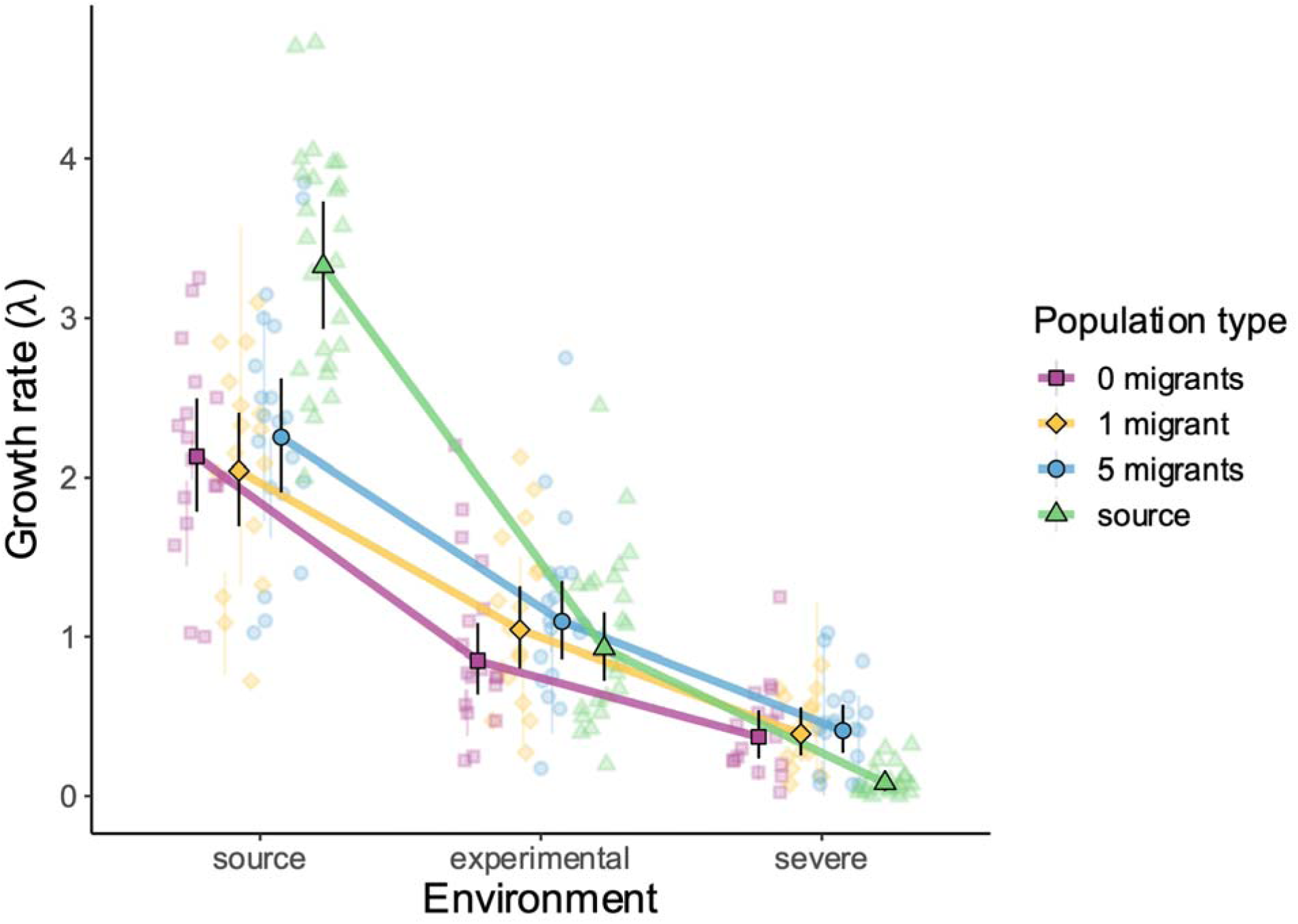
Reciprocal transplant experiment results. Growth rate was measured in three environments: the source environment (standard dry media), the challenging experimental environment, and a severe environment containing a higher concentration of insecticide. Each point outlined in black is the model-estimated mean for each population type (purple squares = 0-migrant populations, yellow diamonds = 1-migrant populations, blue circles = 5-migrant populations, and green triangles = source populations). Black error bars are bootstrapped 95% confidence intervals; for the source populations in the severe environment (green triangles), the black error bars are not visible because they are too small to be seen. Thick lines connecting these points emphasize the treatment-environment interaction. Each point without the black outline represents a single population, with the light error bars representing raw 95% confidence intervals across replicates; points without error bars have no replicates. Points are staggered and jittered to reduce overlap.

Unexpectedly, the experimental and source populations performed similarly in the challenging experimental environment (Figure 5, all confidence intervals of the differences overlap with zero: 95% CI_0-migrant – source_ [-0.404, 0.209]; 95% CI_1-migrant – source_ [-0.220, 0.403]; 95% CI_5-migrant – source_ [-0.153, 0.452]). The experimental populations did not perform as well as expected based on their growth rates at the end of the rescue experiment (Figure 3). Our expectation for growth rates were based on predictions from the linear mixed model created for the rescue experiment with *N*_t-1_ = 40 (the number of parents used in the reciprocal transplant) and gen = 7 (the time closest to the start of the reciprocal transplant). Those predicted values were 1.80, 95% CI [1.32, 2.39], 1.52 [1.12, 2.00], and 1.75 [1.31, 2.26] for the 0-migrant, 1-migrant, and 5-migrant treatments, respectively. The average growth rates we observed in the reciprocal transplant experiment in the experimental environment were on average 40% lower than these estimates, with the means 0.85 [0.64, 1.09], 1.04 [0.80, 1.32], 1.10 [0.86, 1.35].

Furthermore, source populations did not perform as poorly as expected in the experimental environment. We expected them to perform similarly to the first generation of the experimental populations in the rescue experiment prior to adaptation. The prediction given *N*_t-1_ = 40 was 0.63 [0.38, 0.83] for the 0-migrant populations at the start of the rescue experiment, but the mean growth rate of source populations was about 46% higher at 0.92 [0.72, 1.15].

## 4 Discussion

Our study evaluated the effects of different immigration rates on adaptation to a new, challenging environment using experimental populations of *Tribolium castaneum*. We evaluated differences between initially declining populations receiving two immigration rates (one or five migrants per generation) with isolated populations that received no immigration. The migrants were not adapted to the challenging environment, allowing us to examine the degree to which immigration can limit adaptation (Savolainen *et al*., 2013). While the experimental populations declined in size initially, leading some populations to go extinct, around 80% of populations were able to persist for the duration of the seven-generation experiment. Most of these populations had growth rates above the replacement rate and had large population sizes, characteristic of evolutionary rescue. We also found that extinction rate was similar regardless of immigration. Notably, growth rates increased one generation sooner in the isolated populations compared to those with immigration. This delay was brief, as the following generation, growth rates were comparable across the different immigration treatments. We also tested for adaptation using a reciprocal transplant experiment. While some of the results from that experiment were unexpected, overall it provided evidence that our experimental populations were adapted to the insecticide that created the challenging environment. There was also a cost to that adaptation, evident from the experimental populations having lower growth rates than the sources in the benign environment.

### Observed delay in the increase in growth rate: possible mechanisms

Populations without immigration gained a much higher density-independent growth rate (more than doubling on average) compared to the populations with immigration (near or just above the replacement rate) in the fourth experimental generation. The following generation, all populations grew at a similar rate regardless of immigration. The delay we observed may have been due to *i)* differences in density dependence, *ii)* phenotypic plasticity, or *iii)* outcrossing with naive migrants slowing adaptation to the challenging environment.

First, theory shows that reduced negative density dependence in a degraded habitat may enhance rescue (Uecker *et al*., 2014; Czuppon *et al*., 2021). In our system, population densities were low in the experimental populations in the initial generations, and thus negative density-dependent processes were likely relatively weak. The added immigrants increased density slightly relative to the treatment without migrants, and this could have contributed to lower growth rates. However, if this were the case, the populations receiving five migrants each generation should have experienced further reductions in growth than the populations receiving one per generation, as their density was increasing by a greater amount (31% increase for the 5-migrant populations on average compared to a 7% increase for the 1-migrant populations). Thus, density dependence is an unlikely explanation for the one-generation delay in increased growth rates.

Second, growth rate in the challenging environment could be influenced by phenotypic plasticity, which could have led to the lag in increased growth rates. Recently, it has been shown that plasticity in insects can be mediated by associated microbes (Kolodny & Schulenburg, 2020). Variation in the microbial communities in host insects is high (Lange *et al.,* 2023), and the microbial community can evolve, which can facilitate persistence in new and challenging environments (Kolodny & Schulenburg, 2020). Microbes, therefore, can serve as an important source of adaptive phenotypic plasticity (Kolodny & Schulenburg, 2020; Ghalambor *et al.,* 2007). Insect-associated microbes can be transmitted from parent to offspring (Lange *et al.,* 2023), indicating that plasticity could be multi-generational. If the microbial community influences fitness in the presence of deltamethrin, it is possible that plasticity and adaptation in the microbiome helped facilitate the observed increase in growth rate during the rescue experiment. The delay in increase for populations with immigration could be due to the microbiome from the benign environment being continually re-introduced by the migrants.

Third, migration is well known as a constraint to adaptation (Slatkin, 1987). For example, theory has shown that immigration between different habitat types can limit local adaptation through genetic swamping of adaptive alleles (Lenormand, 2002; Bolnick & Nosil, 2007), and reduce the probability (Schiffers *et al*., 2013) or speed (Holt & Barfield, 2015) of evolutionary rescue because of maladaptation in the new environment. It seems likely that migration of naive individuals slowed adaptation to the novel environment, which may have been enhanced by the mating behavior of *T. castaneum*. In this species, males reared in high-quality environments have increased mating rate and insemination success compared to males reared in poor environments (Lewis *et al*., 2012). Thus, male migrants from the benign source environment may have successfully mated with multiple females in the experimental patches, enhancing the rate of gene flow of maladapted alleles. Female migrants would have also likely spread maladapted alleles, as a single female *T. castaneum* can lay over 10 eggs per day (Pray et al., 1949). Thus, regardless of sex, outcrossing with the naive migrants likely contributed to the brief delay in growth rate increase. Given the strength of selection pressure, however, adaptation likely enabled growth rates to increase in the following generation, which we observed on average across all treatments.

We suggest that the increases in population size and growth rate, likely through adaptation and possibly aided by adaptive plasticity, can be described as evolutionary rescue. We further evaluate the effects of adaptation *vs.* plasticity when we discuss the reciprocal transplant experiment below. The finding that rescue can occur with intermediate levels of immigration of not adapted individuals is supported by several modeling studies that assess evolutionary rescue in heterogenous, structured systems (Gomulkiewicz *et al*., 1999; Uecker *et al*., 2014; Tomasini & Peischl, 2020; Czuppon *et al*., 2021). This result has also been shown in a case study of Trinidadian guppies, where rescue was able to occur with gene flow from a maladapted source population (Fitzpatrick *et al.,* 2014, 2020).

### Observed delay in rescue: implications

Our finding of only a brief (one generation) delay of adaptation has practical implications. Such delayed adaptation could be detrimental to populations at imminent risk of extinction due to small size (Gomulkiewicz & Holt, 1995; Melbourne & Hastings, 2008). However, for populations not at immediate risk of extinction, such a delay may be inconsequential, and worth the potential increase in genetic diversity that occurs through immigration. Such increased diversity may prevent inbreeding depression from developing, and improve the ability of the population to adapt to future environmental changes (Frankham *et al.,* 2019).

Thus, these results suggest that naive migrants can be suitable to use in management efforts. Furthermore, they are often more feasible to obtain, either from captive rearing programs (Crone *et al.,* 2007) or from large source populations living in a different habitat (*e.g.,* different host plants in Bisschop *et al.,* 2019) or geographic region (*e.g.,* isolated guppy populations in Fitzpatrick *et al.,* 2020), when habitat matching is not possible.

Additionally, our findings are consistent with refuge strategies to delay evolution of pest resistance to toxins, such as in crops genetically modified to express *Bacillus thuringiensis* (*Bt*). The immigration of susceptible individuals from sections of fields planted in non-*Bt* crops, or growing crops in areas with plentiful alternative host plants for the pest has proven to be highly effective (Carrière *et al*., 2012). In those situations, particularly with 50% or more of a pest population being susceptible, resistance can be successfully managed for multiple generations (Tabashnik *et al*., 2008). Given our much smaller proportion of susceptible individuals, it makes sense that the delay of the evolution of resistance was brief.

### Reciprocal transplant experiment: interpretation of unexpected results

The two most straightforward takeaways from the reciprocal transplant experiment are *i)* the experimental populations had higher growth rates than the source populations in the severe environment, suggesting they had adapted to the insecticide used to create the challenging environment in the rescue experiment and *ii)* the experimental populations had lower growth rates in the benign environment than the source populations, suggesting that there was a cost associated with adaptation to the insecticide. However, the source populations had higher than expected growth rates in the experimental environment, and the experimental populations had lower than expected growth rates in the experimental environment inconsistent with the hypothesis of adaptation. We next discuss *i)* how experimental challenges might have affected the source populations, *ii)* how the common environment generation could have reduced growth rates by eliminating effects of plasticity, and *iii)* how multiple pathways of adaptation in the experimental populations could explain their performance in the reciprocal transplant experiment.

First, the high performance of the source populations in the experimental environment could be due to challenges in implementing the experimental design. For instance, accidental contamination of the source population with deltamethrin, or accidental gene flow from the experimental to the source populations could have occurred during the rescue experiment. We took great care to avoid both forms of contamination, but if contamination nevertheless occurred, it could have allowed the source populations to adapt to low concentrations of the insecticide. The source populations were not adapted to high concentrations of the pesticide, as evidenced by their performance in the severe environment.

A second consideration is that the common environment generation could have masked differences in growth rates, if they were initially caused by plasticity. As described above, microbiome-mediated phenotypic plasticity could have contributed to the increase in growth rate in the rescue experiment (Lange *et al.,* 2023). If microbiome-mediated plasticity drove performance, the generation in a common environment prior to the reciprocal transplant experiment would have alleviated selection pressure and reduced the ability of the experimental populations to tolerate the insecticide. This could help explain the similar performance of the experimental and source populations in the experimental environment. However, there does not appear to be strong evidence for the masking of phenotypic plasticity in the severe or benign environments given that the experimental populations had a much higher growth rate than the sources in the severe environment and a lower growth rate in the benign. We suggest that those differences can be explained by our last point, adaptation, which is discussed below.

Finally, adaptation to deltamethrin likely explains the relative performance of the populations in the three environments. More specifically, it is possible that the experimental populations adapted through more than one mechanism during the rescue experiment. First, adaptation to detoxify the insecticide compound likely occurred, as rapid evolutionary responses to deltamethrin is well documented in pest populations of *T. castaneum* (Collins, 1998; Stuart *et al*., 1998) through the increased expression of cytochrome P450 enzymes (Kalsi & Palli, 2015). In other insect species, this mechanism of resistance to deltamethrin has a confirmed cost (Tchouakui *et al*., 2020), and such a cost is consistent with the reduced performance of experimental populations in the benign environment. The second way that populations could have adapted is by increasing the rate of egg cannibalism to supplement their diets, as cannibalism propensity is a genetically based trait that can evolve (Stevens, 1989) that has been shown to enhance fitness in challenging environments (Via, 1999; Pointer *et al.,* 2021). These potential pathways, evolution of increased detoxification and increased cannibalism, are not mutually exclusive, and both may have facilitated increased population growth in the experimental environment during the rescue experiment. Then, in the reciprocal transplant experiment, the populations were initiated at lower density (40 parents) in the three environments. In this lower density environment, the number of eggs available for cannibalizing was reduced compared to what was available in the latter half of the rescue experiment (in generations 4-7, *N*_t-1_ > 150 on average). Thus, the decreased density reduced the eggs available for cannibalism, limiting one of the adaptive pathways and reducing overall growth rates of the experimental populations in the experimental environment. In the severe environment, growth rates of the experimental populations were low but still greater than the source populations, which can be explained primarily by their evolved ability to detoxify the pesticide.

### Considerations for future experiments

The bottleneck events that occurred during the rescue experiment, some as small as 10 individuals, would likely have longer-term consequences (Frankham *et al*., 1999; Olazcuaga *et al*., 2023). Census population size in a wide range of natural populations is almost always larger than the effective population size (the median *N*_e_/*N*_c_ ratio across plant and animal taxa is approximately 0.1; for *T. castaneum,* it is estimated to be between 0.5-0.9) and bottlenecks of *N*_e_ < 50 can result in reductions in genetic variation and increases in inbreeding (Wade, 1980; Jamieson & Allendorf, 2012). For this reason, we suggest that future work examine the effects of demographic bottlenecks of this severity by propagating experiments like this for longer (*i.e.,* more than 8 generations). Longer-term experiments would be able to test the hypothesis that the benefits of immigration become more apparent over longer time scales. Such experiments would help shed light on the longer-term impacts of immigration for small and isolated populations, an important question when considering whether to implement translocations in declining natural populations (Frankham, 2015, 2016). Genomic methods would also provide important insights for monitoring the genetic contributions of migrants, the genetic differentiation between the experimental and source populations, and for confirming the genes contributing to adaptation (Whiteley *et al*., 2015). Genomics would also allow us to more clearly differentiate between adaptation and plasticity (Koch & Guillaume, 2020).

### Conclusions

This study explored the effects of immigration of naive individuals on large, outbred, and recently diverged populations following population decline due to sudden environmental degradation. To our knowledge, we are the first to experimentally test the effects of the recommended immigration rates of one compared to five migrants per generation on adaptation in a challenging environment. We found that though immigration slowed adaptation in the short-term, extant populations with and without immigration were on average equally well adapted to the challenging conditions after seven generations. Thus, we confirm the findings of previous theoretical work (Tomasini & Peischl, 2020; Czuppon *et al*., 2021) by demonstrating experimentally that evolutionary rescue can occur both with and without immigration following severe environmental change.

## Data Availability Statement

Data and R scripts are publicly available on the Dryad Digital Repository (https://doi.org/10.5061/dryad.79cnp5j3c).

## Conflict of Interest

The authors declare no conflicts of interest.

## Notes

https://doi.org/10.5061/dryad.79cnp5j3c

https://anonymous.4open.science/r/tribolium-rescue-DB19/README.md

### Competing Interest Statement

The authors have declared no competing interest.

### Summary of Updates

Updated Figure 3 and associated statistical analyses.

