## Supplemental Materials for "Immigration delays but does not prevent adaptation following environmental change: experimental evidence"

**Supplementary Materials**

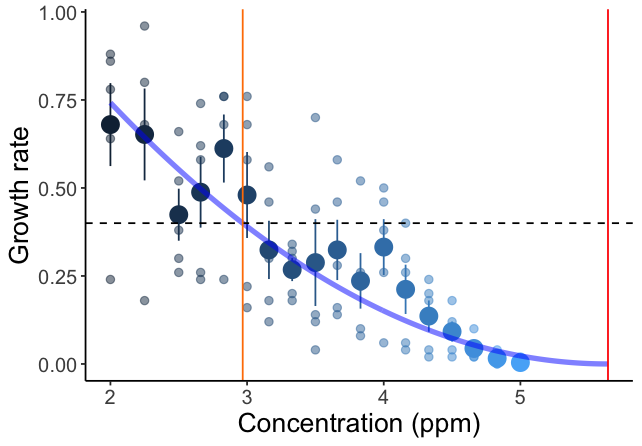

**Figure S1. The dose-response curve** for *Tribolium castaneum* using different concentrations of deltamethrin (DeltaDust, Bayer). The orange line is the experimental concentration chosen based on a goal of growth rate of 0.4 (dashed line). The red line is the concentration used in the severe environment of the reciprocal transplant experiment.

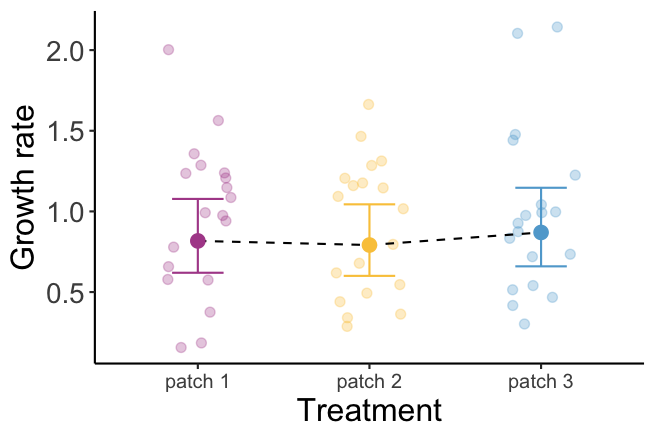

**Figure S2. Order experiment results.** Results of the experiment testing the effects of patch order on growth rate after one generation. Error bars show 95% confidence intervals generated from a linear mixed model including treatment (patch order: position 1-3) as a fixed effect and patch array (19 total across 3 temporal blocks) as a random effect.

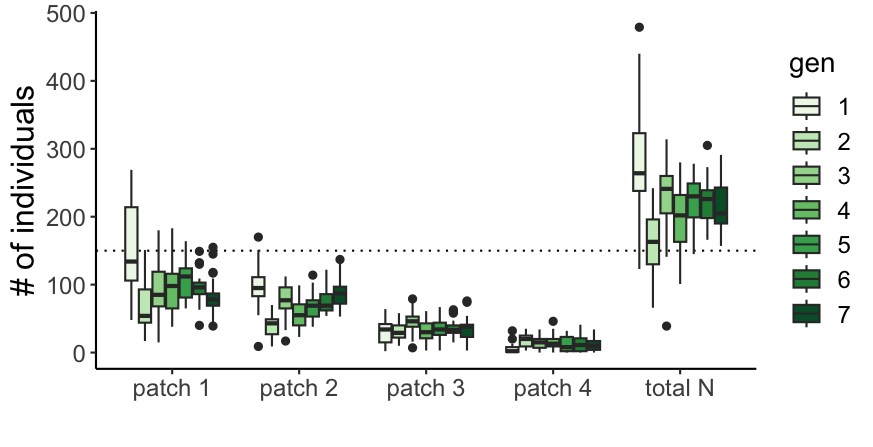

**Figure S3. Source population dispersal array census throughout the rescue experiment.** Distribution of adult beetles among the four patches of the source dispersal arrays (**patch 1-4**) and the total number across all patches (**total N**) for each generation (indicated by darkening shades of green). The dashed line is the initial population size (N = 150) that was used each generation. Box and whisker plots show the median and interquartile range, while whiskers extend to the most extreme data point that is no more than 1.5 times the interquartile range from the box, and points are data more extreme.

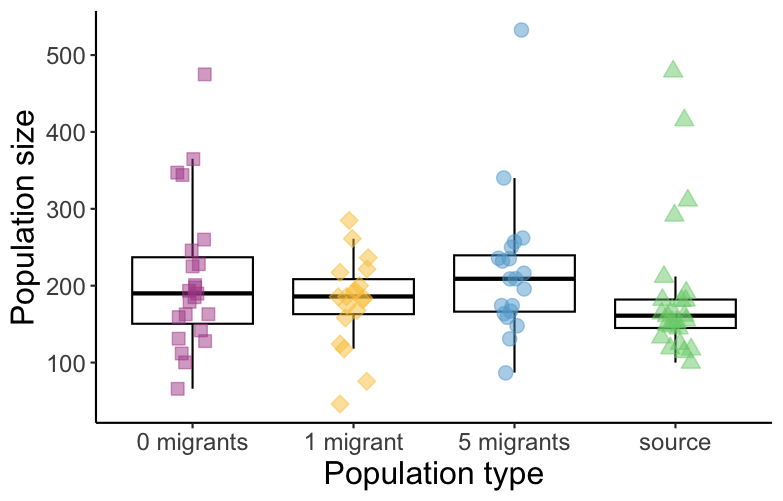

**Figure S4. Common environment generation census sizes.** Total size of each population after the common environment generation (generation 8), in the standard, benign conditions. Each point is the population size for a single population. Box and whisker plots show the median and interquartile range, while whiskers extend to the most extreme data point that is no more than 1.5 times the interquartile range from the box. Between 2-4 replicates were initiated for each population, and some of these populations were allowed to reproduce twice over consecutive days to increase population sizes. We kept the proportion of these double set ups consistent within each treatment. All replicates were initiated with 40 individuals, unless populations were small (*N* < 50) at the conclusion of the rescue experiment; three such populations were initiated with fewer than 40 individuals. The individuals produced in this generation were used to initiate the reciprocal transplant experiment in the subsequent generation. During the reciprocal transplant, populations were split into three groups of 40 individuals, each exposed to a different environment. Additional replication occurred when enough individuals (*i.e.,* 160 or more) were available. In total across all three environments, there were 23-24 replicates of populations from the 0-migrant populations (average replication of 1.1/population/environment), 19-20 replicates (average replication of 1.2/population/environment) from the 1-migrant populations, 22 replicates for the 5-migrant treatment (average replication of 1.1/population/environment), and 24-25 replicates for the source populations (average replication of 1/population/environment).

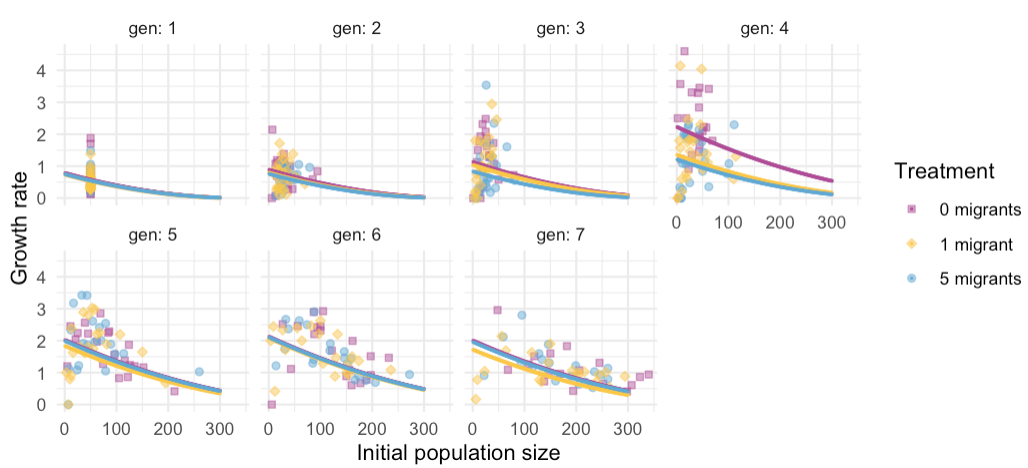

**Figure S5. Growth rate predictions and model fit.** Predictions from the growth rate model across the values of initial population size for each generation during the rescue experiment. The density-independent growth rates were estimated at *N­_0_* = 0, which is the Y-intercept for each generation and treatment combination.

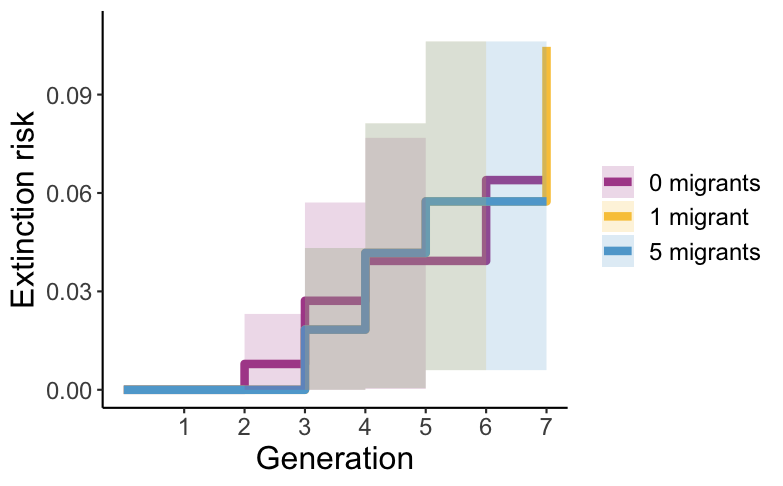

**Figure S6. Extinction risk analysis.** The overall extinction risk for each treatment throughout the rescue experiment using a survival analysis (*ggsurvfit* package in R).

**Table S1. Growth rate comparison through time using different methods.** The density-dependent estimates are predictions from the linear mixed model using average initial population size for each generation and treatment combination, and the density-independent estimates use an initial population size of 0.

| treatment | gen | N | density dependent mean + 95% boot CI | density independent mean + 95% boot CI |
| --- | --- | --- | --- | --- |
| 0 migrants per gen | 1 | 24 | 0.578 (0.351, 0.867) | 0.787 (0.511, 1.137) |
|  | 2 | 24 | 0.746 (0.481, 1.072) | 0.888 (0.601, 1.247) |
|  | 3 | 23 | 0.998 (0.684, 1.371) | 1.130 (0.790, 1.523) |
|  | 4 | 21 | 1.978 (1.519, 2.501) | 2.232 (1.738, 2.804) |
|  | 5 | 20 | 1.452 (1.052, 1.924) | 2.030 (1.533, 2.629) |
|  | 6 | 20 | 1.318 (0.931, 1.769) | 2.131 (1.595, 2.758) |
|  | 7 | 19 | 0.838 (0.540, 1.211) | 2.005 (1.417, 2.709) |
| 1 migrant per gen | 1 | 25 | 0.533 (0.313, 0.818) | 0.736 (0.476, 1.062) |
|  | 2 | 25 | 0.670 (0.416, 0.978) | 0.794 (0.522, 1.134) |
|  | 3 | 25 | 0.924 (0.628, 1.277) | 1.035 (0.725, 1.405) |
|  | 4 | 23 | 1.170 (0.820, 1.580) | 1.356 (0.981, 1.794) |
|  | 5 | 21 | 1.477 (1.085, 1.947) | 1.839 (1.376, 2.380) |
|  | 6 | 20 | 1.410 (1.020, 1.863) | 2.081 (1.574, 2.670) |
|  | 7 | 20 | 0.847 (0.546, 1.207) | 1.716 (1.209, 2.329) |
| 5 migrants per gen | 1 | 25 | 0.544 (0.322, 0.831) | 0.749 (0.480, 1.076) |
|  | 2 | 25 | 0.616 (0.379, 0.912) | 0.760 (0.488, 1.079) |
|  | 3 | 25 | 0.705 (0.452, 1.022) | 0.834 (0.554, 1.164) |
|  | 4 | 23 | 1.009 (0.683, 1.391) | 1.210 (0.853, 1.625) |
|  | 5 | 21 | 1.586 (1.171, 2.059) | 1.995 (1.516, 2.538) |
|  | 6 | 20 | 1.328 (0.943, 1.773) | 2.109 (1.569, 2.738) |
|  | 7 | 20 | 0.959 (0.637, 1.353) | 1.959 (1.416, 2.616) |
